# MiR-34a deficiency enhances nucleic acid sensing and type I IFN signaling in a mouse model of Alzheimer’s disease

**DOI:** 10.1101/2025.09.05.674298

**Authors:** Junling Yang, George Elliot Tsourdinis, Charlotte Holas, Mark Maienschein-Cline, Robert Lalonde, Ken-ichiro Fukuchi

## Abstract

MiR-34a is implicated in aging, cell senescence, inflammation, and neurodegenerative diseases. In order to investigate the role of miR-34a in Alzheimer’s disease (AD), we produced an AD mouse model, Tg-SwDI mice, with whole body/constitutive miR-34a knockout (KO). MiR-34a KO improved long-term memory in Tg-SwDI mice, which was associated with decreases in the ratio of insoluble Aβ42 to Aβ40 and with increases in soluble and insoluble Aβ40 in the cerebral cortex. Anti-Iba1 immunofluorescence revealed increases in activated microglia. Bulk RNA-sequencing of the hippocampus followed by a gene set enrichment analysis (Enrichr) identified “cellular response to type I interferon” and “type I interferon signaling pathway” as the most prominent gene sets in miR-34a KO Tg-SwDI mice compared to miR-34a wild-type Tg-SwDI mice. Many interferon-stimulated genes (ISGs) that characterize interferon responsive microglia (IRM) were upregulated in miR-34a KO Tg-SwDI mice. MiR-34a knockdown strongly enhanced ISGs expression in TLR7 ligand-stimulated BV2 microglia. These results suggest that miR-34a inhibits the transition of microglia to the IRM state that may modulate synaptic and cognitive functions in neurodegenerative diseases and aging.

## 1. Introduction

Alzheimer’s disease (AD) is the most common form of dementia, and hallmark features of its pathology include extracellular amyloid β (Aβ) deposits in the brain parenchyma (amyloid plaques) and intracellular aggregates of hyperphosphorylated tau protein in brain neurons (neurofibrillary tangles, NFTs). Major risk factors for AD such as aging, hypertension, diabetes, obesity, hyperlipidemia, and atherosclerosis, are commonly associated with chronic, low-grade systemic inflammation (1–4). This systemic inflammation contributes to the progression of AD and can trigger microglial activation and subsequent neurodegeneration. Moreover, genetic studies on individuals with late-onset AD have identified many risk genes that are highly expressed in microglia and are involved in inflammatory pathways (5–9).

Microglia play a complex dual role in AD (10–12). On one hand, they protect neurons by engulfing and clearing toxic amyloid-β (Aβ) aggregates and other harmful substances, as well as by secreting neurotrophic factors, including brain-derived neurotrophic factor (BDNF), which support neuronal health (13). On the other hand, microglia can contribute to neurodegeneration by releasing pro-inflammatory and neurotoxic molecules such as tumor necrosis factor-alpha (TNF-α), interleukin-1 beta (IL-1β), reactive oxygen species (ROS), and proteases(14–16). They may also exacerbate damage through excessive synaptic pruning. Understanding the mechanisms that determine this intricate balance between protective and detrimental microglial functions is essential for developing effective therapies for AD.

MicroRNAs (miRNAs) are one class of non-coding endogenous RNAs that post-transcriptionally regulate specific gene expression by several mechanisms and are estimated to control approximately 60% of all mRNAs (17–20). MiR-34a is involved in obesity, diabetes and atherosclerosis, which are high risk factors for AD (21, 22). MiR-34a levels in the brain and blood increase during aging, and increased levels of miR-34a in blood reflect age-dependent brain changes and functional decline in mice (23, 24). MiR-34a levels in the brain are elevated in AD, epileptic seizures, ischemic stroke, and their rodents’ models compared with their controls, and the increased levels of miR-34a in the brain are thought to be involved in pathogenesis (25–28). MiR-34a knockdown by siRNA following cerebral ischemia reperfusion in rats alleviates the tissue injury and neuronal apoptosis by upregulating Notch1 and hypoxia-inducible factor-1α (HIF-1α) (29). Additionally, in a mouse model of transient middle cerebral artery occlusion, miR-34a deficiency reduces blood-brain barrier (BBB) permeability, alleviates disruption of tight junctions, and improves stroke by possibly targeting endothelial cytochrome c-induced apoptosis (30, 31). In cultured microglia, HIV-1 Tat treatment induces a dose- and time-dependent upregulation of miR-34a, leading to microglia activation by directly targeting (downregulating) NLRC5 expression, which negatively regulates NFκB signaling (32). In cultured microglia, p53 activated by reactive oxygen species and DNA damage upregulates miR-34a that targets a transcription regulator, Twist2, leading to downregulation of an anti-inflammatory transcription factor, c-Maf, and microglia activation (33). In cultured microglia, miR-34a decreases phagocytic clearance of Aβ42 by directly targeting/binding TREM2 mRNA (34). Under these conditions, miR-34a appears to be detrimentally augmenting inflammation. In contrast, pre-treatment with miR-34a mimics in rat models of spinal cord injury (SCI) ameliorates microglial activation and neuronal apoptosis by downregulating its targets, Notch1 and Jagged 1, leading to reductions in IL-1β and IL-6 expression (35). MiR-34a knockdown aggravates inflammation and enhances apoptosis in an in vitro model of SCI (36). In LPS-treated RAW264.7 cells (mouse macrophage cell line), miR-34a inhibitor activates NF-kB and increases TNF-α and IL-6 expression by targeting Notch 1 (37). Under the latter conditions, miR-34a seems to be beneficially suppressing inflammation. Thus, miR-34a can exert either pro- or anti-inflammatory effects, as well as neurotoxic or neuroprotective actions, depending on the specific cellular and pathological contexts.

In order to investigate the role of miR-34a in the pathogenesis of AD, we produced an AD mouse model with whole body/constitutive miR-34a knockout (KO or miR-34a^-/-^) and found increases in both soluble and insoluble brain Aβ levels and in activated microglia associated with remarkably increased levels of a full spectrum of interferon-stimulated genes (ISGs) as well as nucleic acid sensing receptor genes.

## 2. Materials and Methods

### 2.1 Experimental animals

Congenic C57BL/6-Tg (Thy1-APPSwDutIowa)BWevn/Mmjax (referred to as Tg-SwDI, Stock No: 34843-JAX) and B6(Cg)-Mir34ATM1LHE/J (referred to as miR-34a KO or miR-34a^-/-^, Stock No: 018279) mice were obtained from Jackson Laboratory. Tg-SwDI and miR-34a KO were repeatedly bred to produce experiment cohorts of miR34a^-/-^ Tg-SwDI and miR-34a^+/+^ Tg-SwDI littermates. Genotyping was performed by PCR using tail biopsy samples, following protocols provided by the Jackson Laboratory. Because Tg-SwDI female mice show more severe AD-like pathological changes than males (38, 39), only female mice were used and analyzed for this study. All animal procedures were prospectively approved by the Institutional Animal Care and Use Committee of the University of Illinois College of Medicine at Peoria. Euthanasia was performed in accordance with the guidelines outlined by American Veterinary Medical Association Panel on Euthanasia.

### 2.2 Behavioral tests

#### 2.2.1 Behavioral schedule and statistics

A battery of behavioral tests was performed on experimental mice at 12 months of age as previously described (40). After measuring body weight and adapting mice to handling, the tests were conducted over a 11-day period as follows: open-field (days 1–3), elevated plus-maze (days 4 and 5), and Morris water maze (days 6–11). In each test, whenever possible, the apparatus was wiped clean with a wet paper towel and dried before the next mouse was introduced to minimize odor cues. Open-field data were analyzed by 2 x 2 x 3 ANOVAs with repeated measures on the day factor. Plus-maze data were analyzed by 2 x 2 x 2 ANOVAs with repeated measures on the day factor. As for Morris water maze, 5 days of testing with repeated measures (2 x 2 x 5 ANOVAs) was used for the 5-day acquisition, and one-way ANOVA and Post-hoc LSD tests were used for the probe trial. Results are expressed as mean ± SD (standard deviation). In all cases, P < 0.05 was considered to be significant.

#### 2.2.2. Exploration and anxiety

Motor activity was measured in the open-field chamber made of white acrylic with a 50 cm × 50 cm surface area. Each mouse was placed in the corner of the open field. The activity in central (25 cm × 25 cm surface area) and peripheral zones was recorded in a 5-min session for 3 consecutive days and analyzed by SMART video tracking software (version 3.0, Harvard Apparatus, Holliston, MA). The distance traveled and the time spent resting (<2 cm/s), moving slow (2–5 cm/s), or moving fast (>5 cm/s) in each zone were measured, as well as the time spent in the periphery and center of the apparatus.

The elevated plus-maze apparatus consisted of 4 arms in a cross-shaped form 70 cm in length with a 10 cm × 10 cm central region. Two of the arms were enclosed on 3 sides by walls (10 cm in height) facing each other, while the other two were open, except for a minimal border (0.5 cm in height) used to minimize falls. Each mouse was placed in the central region, then the number of entries into and the time spent inside enclosed and open arms were measured in a 5-min session on 2 consecutive days with the same video tracking system. The open/total arm entries and duration ratios were also calculated.

#### 2.2.3. Spatial learning and memory

The Morris water maze consisted of a pool of aluminum, 120 cm in diameter with 55 cm high walls (Stoelting Co, Wood Dale, IL), filled with water (22°C) at a height of 31 cm. Powdered milk was dissolved in the water to hide the plexiglass island (escape platform) (10 cm in diameter, 30 cm in height). The milky water was removed every day after a few hours of training and the pool rinsed with clean water. The pool was contained in a room with several external visual cues. The mice were placed next to and facing the wall successively in north (N), east (E), south (S), and west (W) positions, with the escape platform hidden 1 cm below water level in the middle of the SW quadrant. The same video tracking equipment was used to estimate path length and escape latencies in 4-trial sessions for 5 days with a 15-min inter-trial interval. The mice remained on the platform for 5 s. When the mice failed to reach the escape platform within the 1 min cut-off period, they were retrieved from the pool and placed on the platform for 5 s. After swimming, the mice were immediately transferred to a dry plastic holding cage. To maintain body temperature and prevent hypothermia, a red heating lamp was suspended approximately 50 cm above the cages. A cloth towel was placed over half of the cage lid to shield the animal from direct light exposure while still allowing gentle warming if they desire. Mice were kept under these conditions until they were fully dry and active. In the morning after a 5-day acquisition phase, a probe trial was conducted by removing the platform and placing the mouse next to and facing the S side. The time spent in the previously correct quadrant was measured for a single 1-min trial. After the probe trial, the visible platform subtask was conducted, with the escape platform lifted 1 cm above water level and shifted to the NE quadrant. A 17-cm high pole was steadily standing on top of the escape platform as a viewing aid. The same procedure was adopted as with place learning except that the subtest was conducted on a single day.

### 2.3 Immunohistochemical analyses

After completion of behavioral testing, mice were deeply anesthetized with a mixture of ketamine (200 mg/kg) and xylazine (20 mg/kg), followed by transcardial perfusion with cold PBS. Brains were rapidly removed; the left hemisphere’s neocortex and hippocampus were dissected and stored at −80 °C for molecular analyses. The right hemisphere was fixed in 4% paraformaldehyde for 48 h, cryoprotected in 30% sucrose in 0.1 M PBS for 48 h at 4 °C, embedded in OCT compound (Tissue-Tek), and frozen. Coronal sections (35 µm) were collected on a cryostat from approximately 1.3 to 3.4 mm posterior to bregma. For immunostaining, three sections per mouse were selected at ∼600 µm intervals. To enhance the detection of Aβ deposits, a mixture of anti-Aβ antibodies — 6E10, which targets amino acids 1-16 of the Aβ peptide, and 4G8, which binds to amino acids 17-24 —was used as reported previous (41). Antigen retrieval was performed in antigen retrieval buffer (Cat# ab64236, Abcam) at 85 °C for 10 min, followed by cooling to room temperature. Sections were washed in 0.1 M Tris buffer (TB), then treated with 1% hydrogen peroxide in 10% methanol (in TB) for 1 h to quench endogenous peroxidase activity. After sequential washes in TB and TBS (pH 7.5), blocking was carried out with M.O.M.™ reagent (Cat# BMK-2202, Vector Laboratories) for 1 h, followed by a brief TBS wash. Sections were incubated for 48 h at 4 °C with 6E10 (1:1000; BioLend Cat# 803001, RRID: AB_2564653) and 4G8 (1:1000; BioLegend Cat# 800701, RRID: AB_2564633) diluted in M.O.M.™ diluent. After rinsing (4 × 15 min in TBS), sections were incubated with a biotinylated anti-mouse IgG secondary antibody (Vector Laboratories) diluted in 1% BSA in TBS-T for 2 h at room temperature. Detection was performed using the avidin-biotin complex (ABC) method (Vector Laboratories) with 3,3′-diaminobenzidine (DAB; Sigma-Aldrich, St. Louis, MO) as the chromogen. Sections were washed in water, dehydrated through graded ethanol, cleared in xylene, and cover slipped using mounting medium. Images were obtained using a Zeiss Axioscan. For microglial staining, sections underwent the same antigen retrieval procedure as above, then were incubated with rabbit anti-IBA1 (1:1000; FUJIFILM Wako Pure Chemical Corporation Cat# 019-19741, RRID: AB_839504) for 48 h at 4 °C. After rinsing (4 × 15 min in TBS), sections were incubated with Alexa Fluor-conjugated anti-rabbit IgG (1:500; Cell Signaling Technology Cat# 4412, RRID: AB_1904025) for 2 h at room temperature in the dark. Nuclei were counterstained with DAPI, and sections were mounted using Shandon™ Immu-Mount™ (Thermo Fisher Scientific). Images were acquired using an Olympus FV3000 confocal microscope (Olympus America Inc.) with consistent settings using 40× objectives. Dentate gyrus of hippocampus was analyzed by Image Pro 10 image analysis software (Media Cybernetics, Silver Spring, MD) capable of color segmentation and automation via programmable macros. Data was expressed as mean ± SD).

### 2.4 Quantification of brain Aβ by ELISA

The left neocortex, previously stored at -80°C, was lysed using the Bio-Plex cell lysis kit (Bio-Rad Laboratories, Hercules, CA) and homogenized with a Bullet Blender STORM (Laboratory Supply Network, Atkinson, NH) following the manufacturer’s instructions. Samples were centrifuged at 12,000 × g for 10 min at 4°C, and the resulting supernatants, containing buffer-soluble Aβ, were collected. Protein concentrations in the supernatants were determined using the Bio-Rad protein assay (Bio-Rad Laboratories). The remaining pellets, containing insoluble Aβ, were further homogenized in 5M guanidine hydrochloride using Bullet Blender STORM and then rock-shaken for 3-4 h at room temperature. Levels of buffer soluble and insoluble Aβ in the neocortex were quantified by Aβ42 and Aβ40 ELISA kits (Invitrogen, Carlsbad, CA) according to the manufacturer’s protocol.

### 2.5 Western blot

Protein lysates were used to assess IFITM3 and CSF1R levels by western blotting. After quantifying protein concentrations, samples were separated under reducing conditions on 10% SDS–PAGE gels and transferred to PVDF membranes. Membranes were incubated overnight at 4°C with primary antibodies against IFITM3 (1:1000; Abcam Cat# ab15592, RRID: AB_2122095), CSF1R (1:1000; R and D Systems Cat# AF3818, RRID: AB_884158), and GAPDH (1:5000; Millipore Cat# MAB374, RRID: AB_2107445). After three washes in PBST, membranes were incubated with species-appropriate HRP-conjugated secondary antibodies: anti-rabbit (1:2000; Cell Signaling Technology Cat# 7074, RRID: AB_2099233), anti-sheep (1:1000; R and D Systems Cat# HAF016, RRID: AB_562591), and anti-mouse (1:3000; Cell Signaling Technology Cat# 7076, RRID: AB_330924). Protein bands were detected using enhanced chemiluminescence (Amersham, Arlington Heights, IL). Band intensities were quantified by densitometric analysis using an HP Scanjet G3010 and ImageJ v1.40 (NIH).

### 2.6 RNA sequencing and bioinformatics analysis

Total RNA was extracted from the left hippocampus using the Direct-zolTM RNA kit (Zymo Research Corporation, Irvine, CA). RNA sequencing was performed on DNBSEQ high-throughput platform at BGI Tech Solution NGS lab (Hong Kong, CHN). Raw reads were aligned to the mouse reference genome mm10 using STAR (42). ENSEMBL genes were quantified using FeatureCounts as raw read counts (43). Differential expression statistics (fold-change and p-value) were computed using edgeR using generalized linear models to compute differential expression due to genotype and treatment in a two-factor design, including an interaction term (44, 45). Pairwise comparisons between groups were also computed using the “exactTest” function. Gene expression was normalized using TMM factors in edgeR. In all cases, p-values were adjusted for multiple testing using the false discovery rate (FDR) correction of Benjamini and Hochberg (46). PCA plots were generated from log-scaled normalized expressions. We identified gene expression patterns with dependent effects of genotype and treatment through a clustering analysis. Differentially expressed genes (DEGs) for clustering were selected based on interaction term FDR < 0.2. Differential profiles of these genes were computed as the product of the log fold-change and -log p-value for each pairwise comparison and normalized by dividing by their standard deviation. We then performed k-means clustering on these profiles with ten random initializations on a range of cluster numbers k (2 to 20). We determined the optimal k by evaluating clustering reproducibility statistics for each set of results, similar to the approach outlined by Senbabaoglu and co-workers: for each k, we computed the pair-wise distance between clustering results as the fraction of co-clustered feature pairs across two clustering results relative to the number of co-clustered feature pairs within each result individually (47). This difference was averaged across all result pairs for each value of k, and the largest k with an average distance less than 1e-5 was selected as the ideal clustering granularity –k = 4 was determined to be the best number of clusters. DEGs were entered into Enrichr database (Enrichr, https://maayanlab.cloud/Enrichr/) with threshold of log2foldchange (FC) < -0.1 or > 0.1 and FDR < 0.05 (48, 49).

### 2.7 Validation of miR-34a knockdown and quantitative PCR analysis of type I interferon signaling in gardiquimod-treated BV2 cells

BV2 microglial cells were seeded in 6-well plates at a density of 4.8 x 10^5^ cells/ml and cultured overnight in DMEM (ATCC formulation) supplemented with 10% fetal bovine serum (FBS) under standard conditions. Cells were transfected with 25 nM of either a miR-34a inhibitor or a negative control miRNA using Lipofectamine RNAiMAX Transfection Reagent (Life Technologies, Grand Island, NY) according to the manufacturer’s instructions. Twenty-four hours after transfection, cells were treated with 0.2 μg/mL Gardiquimod (Cat# tlrl-gdqs-10, InvivoGen, San Diego, CA), a TLR7 agonist known to stimulate type I interferon responses. After 24 hours of Gardiquimod treatment, total RNA was extracted using the Direct-zol™ RNA Kit (Zymo Research, Irvine, CA), and RNA concentration and purity were assessed using a NanoDrop spectrophotometer. cDNA was synthesized using a high-capacity cDNA reverse transcription kit (Bio-Rad Laboratories) following the manufacturer’s protocol.

Quantitative PCR (qPCR) was performed using SYBR Green Master Mix (Bio-Rad Laboratories) on a CFX-384 Real-Time PCR System (Bio-Rad Laboratories). Primer sequences are listed in Table 1. MiR-34a expression was assessed using microRNA assays (Qiagen) with RNU1A1 as internal controls. Relative gene expression levels were calculated using the 2^−ΔΔCt^ method.

**Table 1.**
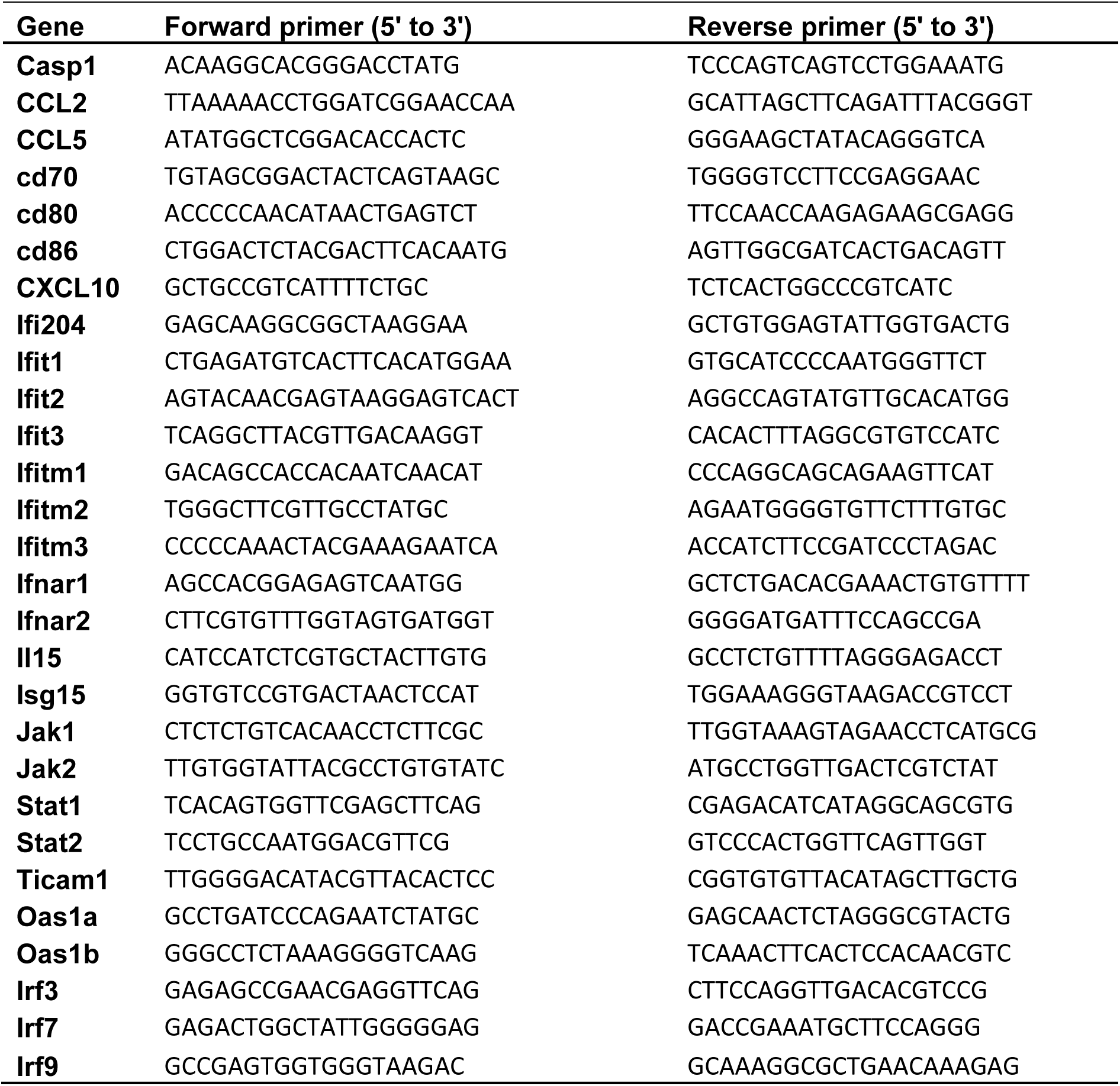
Primer list of interferon stimulated genes studied for qPCR.

### 2.8 Statistic analysis

Statistical analyses were performed using SPSS version 19. Intergroup differences were evaluated using repeated measures ANOVA and two-tailed t-tests were used for comparisons between two independent groups. P < 0.05 was considered statistically significant.

## 3. Results

### 3.1 MiR-34a deficiency enhances exploratory activities and spatial memory in mice

#### 3.1.1 Exploration and anxiety

In the mouse open-field test (Figure 1), day effects were observed in peripheral time (F(2,78)=26.117, P=0.000) and resting time (F(2,78)=100.961, P=0.000), which increased across the days and in center time (F(2,78)=26.073, P=0.000), moving fast time (F(2,78)=95.442, P=0.000), moving slow time (F(2,78)=32.365, P=0.000), and travelled distance (F(2,78)=104.881, P=0.000), which decreased across days, indicating habituation of exploratory activity over time. The miR-34a KO groups had less peripheral time (F(1,39)=8.813, P=0.005), more center time (F(1,39)=8.814, P=0.005), more moving slow time (F(1,39)=53.496, P=0.000), and less resting time (F(1,39)=26.099, P=0.001) than the miR-34a WT groups, suggesting that miR-34a KO decreased anxiety in mice. The miR-34a KO x day interaction (F(2,78)=4.879, P=0.012) in moving fast time, moving slow time (F(2,78)=5.248, P=0.011) and resting time (F(2,78)=9.443, P=0.000) were caused by larger KO effects on days 2 and 3 than the initial test day. There is a trend of miR34a KO x APPSwDI transgene x day interaction in moving fast (F(2,78) =2.986, P=0.061), moving slow (F(2,78)=2.642, P=0.088), resting time (F(2,78)=3.235, P=0.052), and travelled distance (F(2,78)=3.035, P=0.057). These results indicate that miR-34a KO mice are more active than miR-34a WT mice across the days.

**Figure 1.**
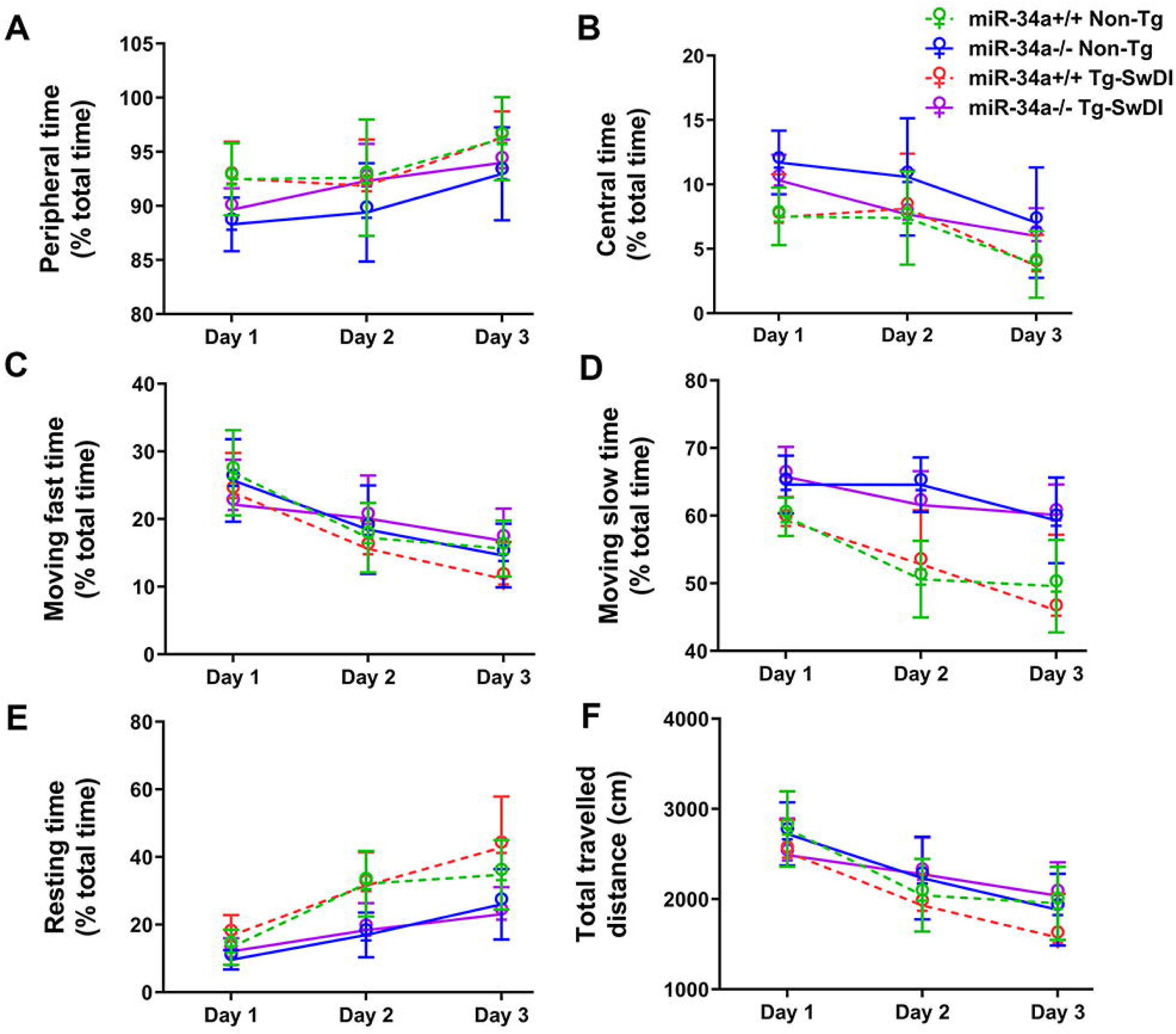
Effects of miR-34a KO (miR-34a^-/-^) on exploratory activity and anxiety-like behaviors in open-field. The percentages of peripheral time (A), central time (B), moving fast time (C), moving slow time (D), resting time (E), and total travelled distances (F) in a 5-min session are shown as mean ± SD for 3 days. (n=9-12/group).

In the elevated plus-maze test (Table 2), there were day effects in open arm entries (F(1,39)=123.376, P=0.000), open arm duration (F(1,39)=334.804, P=0.000), enclosed arm entries (F=(1,39)=7.413, P=0.01), open/total entries (F(1,39)=80.321, P=0.000), and open/total duration (F=(1,39)=348.383, P=0.000), which decreased across days and enclosed arm duration (F(1,39)=238.477, P=0.000), which increased across days, as a result of safer exploration of the maze over time. Tg-SwDI mice had more open arm entries (F(1,39)=5.404, P=0.025), open arm duration (F(1,39)=9.475, P=0.004), open/total entries (F(1,39)=16.255, P=0.000), and open/total duration (F(1,39)=9.622, P=0.004) but less enclosed arm duration (F(1,39)=8.175, P=0.007) than Non-Tg mice, indicating riskier exploration of the maze on the part of the transgenic groups. These effects are amplified in the KO group relative to the WT group as indicated by significant transgene x miR34a interactions in open arm entries (F(1,39)=5.292, P=0.027), open arm duration (F(1,39)=6.802, P=0.013), enclosed arm duration (F(1,39)=6.800, P=0.013), and open/total duration (F(1,39)=6.932, P=0.012). The transgene x miR34a x day interactions in open arm entries (F(1,39)=5.793, P=0.021) and enclosed arm entries (F(1,39)=4.258, P=0.046) are due to larger effects on day 2 than day 1.

**Table 2.** Elevated Plus. Effects of miR-34a KO on exploratory activity in elevated plus-maze (n=9-12, mean ± SD; * P < 0.05).

| Measures | miR-34a<sup>+/+</sup> Non-Tg | miR-34a<sup>-/-</sup> Non-Tg | miR-34a<sup>+/+</sup> Tg-SwDI | miR-34a<sup>-/-</sup> Tg-SwDI |
| --- | --- | --- | --- | --- |
| **Day 1** |  |  |  |  |
| **Open arms** |  |  |  |  |
| Entries | 14.45 ± 3.80 | 13.17 ± 4.73 | 15.40 ± 4.17 | 14.90 ± 4.07 |
| Duration (s) * | 96.70 ± 27.18 | 83.09 ± 26.85 | 100.16 ± 18.17 | 120.57 ± 46.79 |
| **Enclosed arms** |  |  |  |  |
| Entries | 14.55 ± 3.30 | 12.25 ± 2.67 | 12.40 ± 5.38 | 10.10 ± 4.41 |
| Duration (s) | 162.35 ± 25.93 | 178.81 ± 32.51 | 157.83 ± 29.50 | 140.30 ± 46.91 |
| **Open/total ratio (%)** |  |  |  |  |
| Entries | 25.02 ± 3.98 | 25.22 ± 5.90 | 28.59 ± 4.75 | 30.72 ± 6.05 |
| Duration | 32.23 ± 9.06 | 27.70 ± 8.95 | 33.39 ± 6.06 | 40.52 ± 15.60 |
| **Day 2** |  |  |  |  |
| **Open arms** |  |  |  |  |
| Entries * | 6.00 ± 3.66 | 3.50 ± 1.78 | 5.10 ± 3.34 | 10.50 ± 5.28 |
| Duration (s) * | 30.66 ± 17.24 | 23.81 ± 10.14 | 34.48 ± 18.99 | 74.45 ± 38.66 |
| **Enclosed arms** |  |  |  |  |
| Entries | 12.73 ± 4.96 | 8.08 ± 2.02 | 9.90 ± 6.01 | 11.00 ± 4.64 |
| Duration (s) * | 236.61 ± 29.45 | 248.44 ± 16.09 | 236.21 ± 35.30 | 180.15 ± 53.59 |
| **Open/total ratio (%)** |  |  |  |  |
| Entries | 15.27 ± 4.88 | 15.09 ± 4.56 | 18.52 ± 7.90 | 24.54 ± 5.41 |
| Duration * | 10.22 ± 5.75 | 7.94 ± 3.38 | 11.49 ± 6.33 | 24.82 ± 12.89 |

#### 3.1.2 Spatial learning and memory

The results of the Morris water maze test are shown in Figure 2. In the acquisition phase, significant day effects were observed for both path lengths (F(4,156)=36.556, P=0.000) and escape latencies (F(4,156)=33.263, P=0.000), indicating learning over time. Nevertheless, Tg-SwDI mice had longer latencies than Non-Tg mice (F(1,39)=5.693, P=0.022, Figure 2B), suggesting that the APPSwDI transgene decreased learning ability in mice. In addition, there are APPSwDI transgene x miR34a x day interactions in path lengths (F(4,156)=2.662, P=0.041, Figure 2A) and latencies (F(4,156)=2.863, P=0.029, Figure 2B) ), mostly due to slower acquisition on the part of miR-34a KO Tg-SwDI mice on days 2 to 4. But during the probe trial, the percentage of time spent in the correct quadrant for miR-34a KO mice was greater than that of miR-34a WT mice (F(3, 39) =7.178, P=0.011, Figure 2B) regardless of transgenic status. On the contrary, no intergroup differences occurred when the platform became visible (MiR-34a^+/+^ Non-Tg: 392.14 ± 200.39 cm, 25.49 ± 12.32 seconds; miR-34a^-/-^ Non-Tg: 464.14 ± 241.87 cm and 34.19 ± 22.45 seconds; miR-34a^+/+^ Tg-SwDI: 466.85 ± 158.05 cm and 32.92 ± 10.63 seconds; miR-34a^-/-^ Tg-SwDI: 480.45 ± 298.55 cm and 42.79 ± 34.56 seconds, shown as path lengths and latencies, respectively). These results indicate that miR-34a KO improves spatial long-term memory in mice and Tg-SwDI mice show learning deficits in the Morris water maze at 12 to 13 months of age.

**Figure 2.**
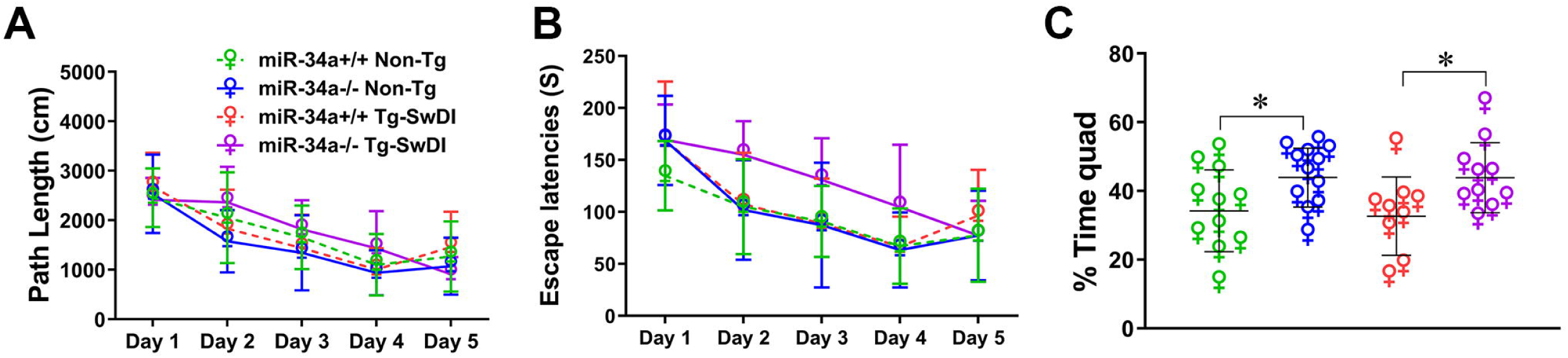
Effects of miR-34a KO (miR-34a^-/-^) on cognitive function in Morris water maze. Total path lengths (cm) per day (A), Total escape latencies per day (s) (B), and Percentages of time spent in the target quadrant in which the hidden platform were previously placed are shown as mean ± SD (C). (n=9-12/group)

### 3.2 MiR-34a KO decreased the ratio of insoluble Aβ42 to Aβ40 in cerebral cortex in Tg-SwDI mice

Tg-SwDI mice primarily exhibit amyloid deposition in the subiculum and dentate gyrus (50). To determine if miR-34a KO influences Aβ pathology, we evaluated Aβ deposition in the dentate gyrus of 13-month-old Tg-SwDI mice by immunohistochemistry using 6E10 and 4G8 antibodies (Figure 3A and B). The Aβ loads were indicated by average percentages of area showing Aβ immunoreactivity in the polymorph layer of the dentate gyrus (Figure 3C). No difference was found in Aβ loads by immunohistochemistry between miR-34a KO and WT Tg-SwDI mice.

**Figure 3.**
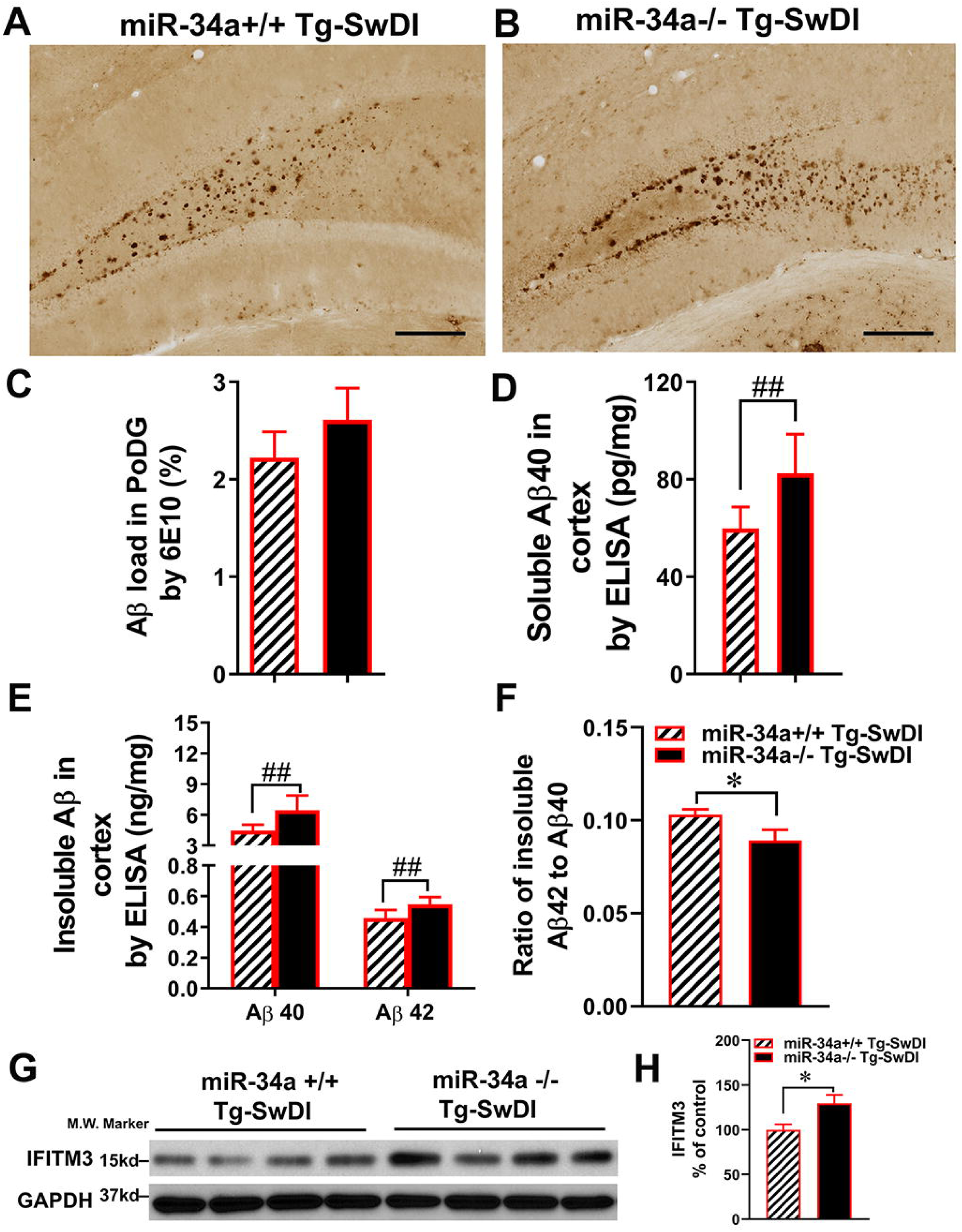
Effects of miR-34a KO (miR-34a^-/-^) on Aβ load and APP processing in the brain. Detection of Aβ deposits by immunohistochemistry with anti-Aβ antibody, 6E10 in miR-34a^+/+^ (A) and miR-34a^-/-^ (B) Tg-SwDI mice. Scale bars 200 µm. Average percentages of areas showing Aβ immunereactivity measured by morphometry in the polymorph layer of the dentate gyrus (PoDG) are shown (C). The levels of soluble (D) and insoluble (E) Aβ40 and Aβ42 in the cerebral cortex were measured by ELISA. Data are shown as mean ± SD. The ratios of insoluble Aβ42 to Aβ40 in the cerebral cortex (F). Effects of miR-34a KO on levels of IFITM3 (G and H) in lysates of the cerebral cortex were analyzed by western blotting using anti-ITIFM3 antibody (G), and bar graph represents the results of densitometric analysis of IFITM3 (H) after normalizing with GAPDH levels. (A-F: n=9-10/group, *P < 0.05, #P < 0.01 and ##P < 0.001; G and H: n=4, *P < 0.05).

The amounts of buffer soluble and insoluble Aβ40 and Aβ42 in the cerebral cortex were determined by ELISA. MiR-34a KO significantly increased soluble Aβ40 in the cortex in Tg-SwDI mice (59.87 ± 8.78 vs 82.45 ± 16.14 pg/mg protein, P < 0.001, Figure 3D). Insoluble Aβ40 and Aβ42 significantly increased in the cortex in miR-34a KO Tg-SwDI mice (6.44 ± 1.14 and 0.55 ± 0.05 ng/mg wet weight, respectively) compared with miR-34a WT Tg-SwDI mice (4.47 ± 0.58 and 0.46 ± 0.05 ng/mg wet weight, respectively, P < 0.001 for both Aβ40 and Aβ42, Figure 3E). It has been shown that the higher ratio of Aβ42/Aβ40 drives Aβ mixtures into neurotoxic oligomeric assemblies (51, 52). Because Aβ42 in the buffer-soluble fraction of Tg-SwDI mice at 13 months of age is undetectable by ELISA, we determined the Aβ42/Aβ40 ratio of buffer-insoluble Aβ in the cortex of Tg-SwDI mice. The cortical insoluble Aβ42/Aβ40 ratio in miR-34aKO mice (0.089 ± 0.018) is significantly lower than that (0.103 ± 0.009) in miR-34a WT Tg-SwDI mice (Figure 3F, p < 0.05).

Because miR-34a KO increased levels of cerebral Aβ as well as mRNA levels of IFITM3 (See below) in Tg-SwDI mice and because IFITM3 can enhance γ-secretase activity, expression levels of IFITM3 in the cerebral cortex were analyzed by western blotting using anti-IFITM3 antibody (53). The protein expression levels of IFITM3 in the cerebral cortex in miR-34a KO Tg-SwDI mice were significantly higher than those in miR-34a WT Tg-SwDI mice (Figure 3G and H, P < 0.05).

### 3.3 MiR-34a KO enhanced microglial activation in Tg-SwDI mice

Because Tg-SwDI mice develop Aβ deposition predominantly in the subiculum and dentate gyrus (see Figure 3A, also) (50), we further investigated glial activations in the dentate gyrus of mice (Figure 4). IBA1 immunostaining shows that an increased cell body-to cell size ratio is associated with enhanced microglia activation (54). MiR-34a KO significantly increased the total number of IBA1-immunoreactive microglia in the dentate gyrus in Tg-SwDI mice (Figure 4A-C, P < 0.001, Scale bar: 50µm), as well as microglial cell body to cell size ratios by morphometric analysis (Figure 4D, P < 0.001).

**Figure 4.**
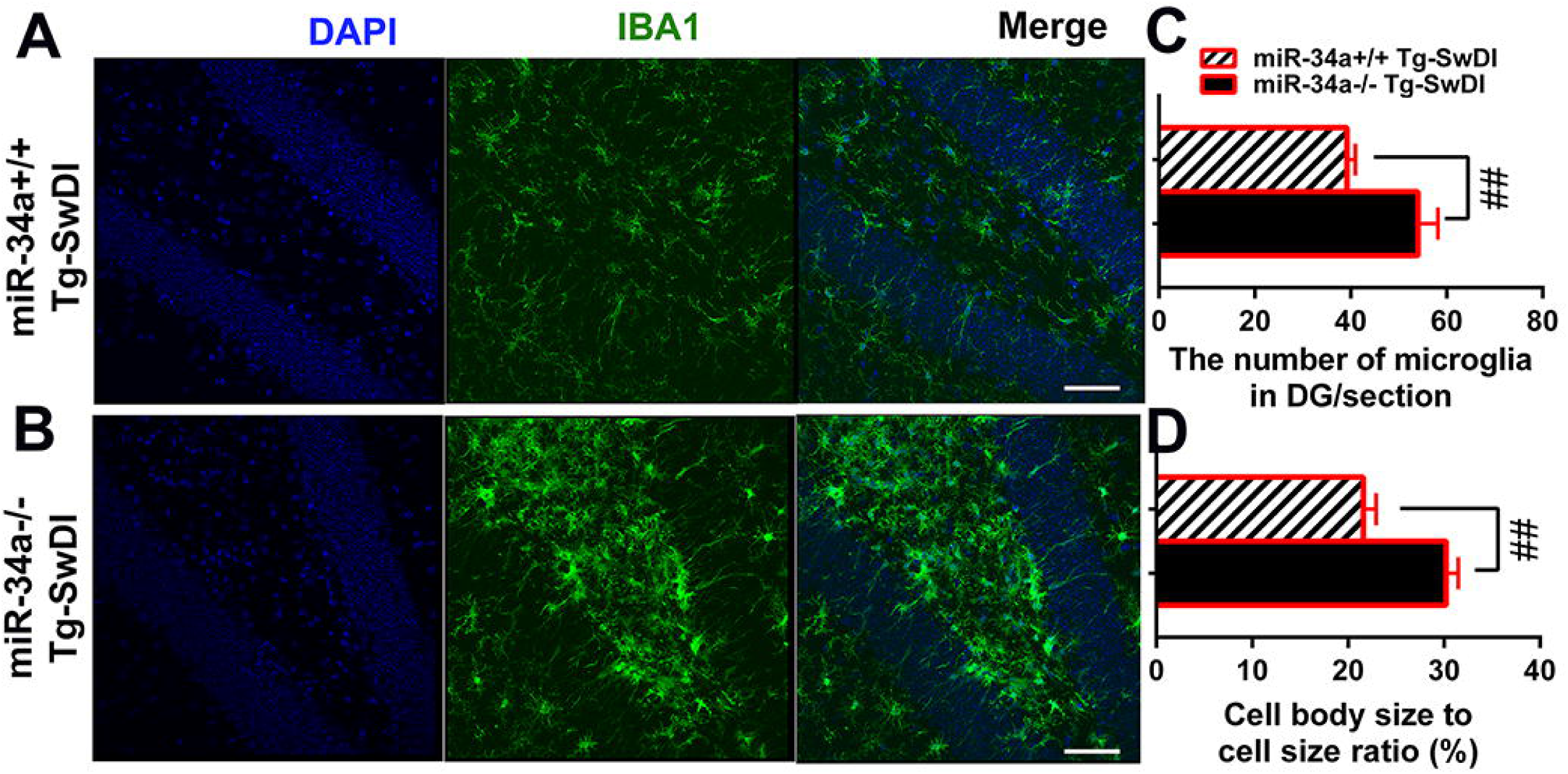
Effect of miR-34a KO on microglia activation by IBA1 staining. Brain sections were subjected to immunofluorescence staining using anti-IBA1 antibody (green) and DAPI staining (blue). The dentate gyrus from miR-34a^+/+^ (A) and miR-34a^-/-^ (B) Tg-SwDI mice are shown. Scale bars 50 µm. The total number of microglia/35 µm thick section (C) and the ratio of microglial cell body to cell size (D) in the dentate gyrus were determined. (n=9-10/group, ##P < 0.001).

To elucidate the role of miR-34a in hippocampal transcriptomics, we conducted bulk RNA sequencing of hippocampal tissue isolated from mice. We found that miR-34a KO significantly increased expression levels of many disease-associated markers (DAM) including Apoe, Tyrobp, B2m, Ctsb, Trem2, Clec7a, Axl, and Csf1 (Figure 5A, *Q < 0.05, ##Q < 0.001), suggesting that miR-34a KO increased activated microglia with expression profiles of DAM in Tg-SwDI mice. Additionally, miR-34a KO increased markers of homeostatic microglia, including C1q and CSF1R (Figure 5A, *Q < 0.05, and ##Q < 0.001). Because CSF1R signaling stimulates microglial proliferation and survival and because CSF1R is an experimentally validated target of miR-34a (Figure 5B) (55–57), we determined protein expression levels of cerebral CSF1R by western blotting in miR-34a KO and miR-34a WT Tg-SwDI mice. There was a large increase in CSF1R in miR-34a KO Tg-SwDI mice compared with miR-34a WT Tg-SwDI mice (Figure 5C, P < 0.001), suggesting that increased CSF1R signaling in miR-34a KO mice may induce microglia proliferation and activation in response to Aβ accumulation.

**Figure 5.**
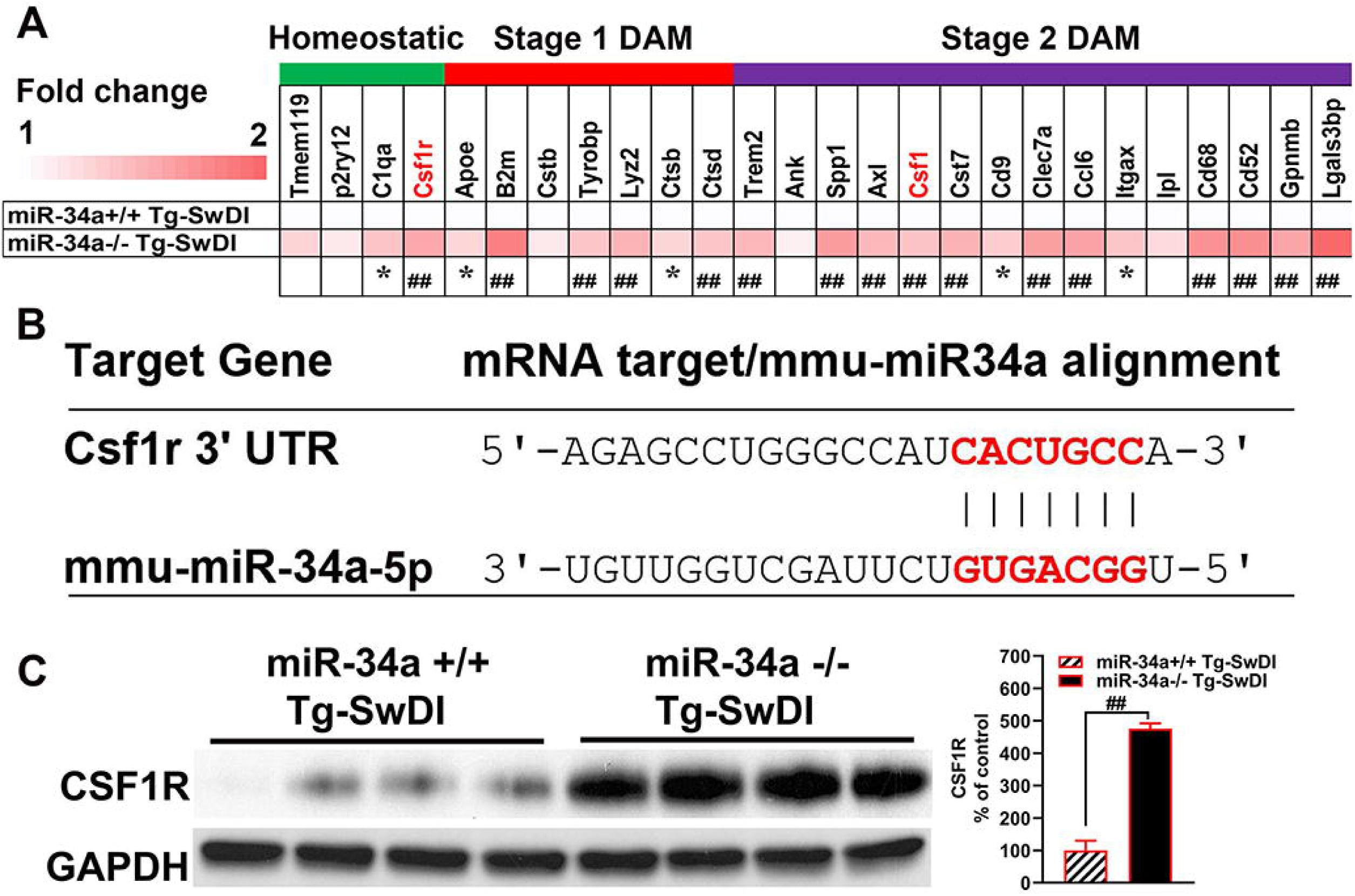
Effects of miR-34a KO on expression levels of homeostatic and DAM genes. Expression levels (Heatmap) of marker genes for three different microglial stages (homeostatic, stage 1 DAM, and stage 2 DAM) are shown as fold changes miR-34a^-/-^ to miR-34a^+/+^ Tg-SwDI mice, based on bulk RNAseq of the hippocampus (A). (n=5/group, *Q < 0.05, ##Q < 0.001) Sequence alignment of the mature miR-34a with CSF1R 3’ untranslated region (3’UTR). The seed sequences and target mRNA are highlighted in red (B). Effects of miR-34a KO on levels of CSF1R in lysates of the cerebral cortex were analyzed by western blotting using CSF1R antibody (C) and bar graph represents the results of densitometric analysis of CSF1R after normalizing with GAPDH levels. (n=4/group, ##P < 0.001).

### 3.4 MiR-34a KO activated type I interferon signaling in Tg-SwDI mice revealed by RNA-seq analysis

Compared with their Non-Tg littermates, the numbers of differentially expressed genes (DEGs) with the false discovery rate (FDR) < 0.05 cut off are 2,178 and 796 for miR-34a KO and miR-34a WT Tg-SwDI mice, respectively (Supplementary Table 1). Thus, miR-34a KO almost tripled the number of DEGs in Tg-SwDI mice. Of the 2,178 DEGs in miR-34a KO Tg-SwDI mice, 1,684 were up-regulated and 494 were down-regulated, and of the 796 DEGs in miR-34a WT Tg-SwDI mice, 677 were upregulated and 119 were down-regulated. When miR-34a KO Tg-SwDI mice were compared with miR-34a WT Tg-SwDI mice, 268 genes were found to be DEGs (FDR < 0.05) with 252 up- and 16 down-regulated genes. Compared with miR-34a WT Non-Tg mice, a total of 108 genes in miR-34a KO Non-Tg mice were identified as DEGs with 21 up-and 87 down-regulated. Consequently, miR-34a KO increases the number and expression levels of up-regulated genes in Tg-SwDI mice but reverses the number and expression levels of up-regulated genes in Non-Tg littermates. These results indicate that hippocampal miR-34a regulates expression of a small number of genes under normal conditions but regulates expression of a large number of genes in response to AD-like brain changes. This suggests that miR-34a may play an important role in modulating hippocampal gene expression and, ultimately, phenotypic manifestation in AD. DEGs were subjected to the Enrichr database. When a paired-wise comparison was made between miR-34a KO Tg-SwDI and miR-34a WT Tg-SwDI mice, type I interferon signaling pathway and cellular response to type I interferon were ranked at top 1 sets based on GO-BP (gene ontology biological process) (Figure 6A, Supplementary Table 2). Activation of the type I IFN pathway in miR-34a KO Tg-SwDI was clearly highlighted by the volcano plot based on DEGs compared to miR-34a WT Tg-SwDI mice (Figure 6B). Thus, miR-34a KO induced type I interferon (IFN) signaling in Tg-SwDI mice with significant Q-values (Supplementary Table 3). The upregulated DEGs include Ifnar2, Stat2, Oas1a, Oas2, Oas3, Irf7, Gbp2, Isg15, and Irf9, which are commonly involved in NOD-like receptor signaling, double-stranded RNA binding, and RIG-I-like receptor signaling pathways. Because these signaling pathways and the volcano plot (Figure 6B) include nucleic acid sensors and because activation of nucleic acid sensors leads to induction of the type I interferon signaling, we compared expression levels of nucleic acid sensors between miR-34a KO and WT Tg-SwDI mice. Expression levels of nucleic acid sensors including Tlr3, Ifih1/MDA5, Ddx58/RIG-1, Ifi204, Ifi209, Ifi211, and Zbp1 in miR-34a KO Tg-SwDI mice are significantly higher than those in miR-34a WT Tg-SwDI mice (Supplementary Table 1). Importantly, many upregulated genes involved in type I interferon signaling as well as nucleic acid sensors in miR-34a KO Tg-SwDI mice can be targeted by miR-34a and are summarized in Supplementary Table 3, suggesting that miR-34a KO may contribute to activation of type I interferon signaling through upregulation of these potential target genes and nucleic acid sensors. Additionally, almost all genes of the core IRM gene signature (58) were upregulated in miR-34a KO Tg-SwDI mice (Supplementary Table 3), suggesting that miR-34a KO induced IRMs in Tg-SwDI mice.

**Figure 6.**
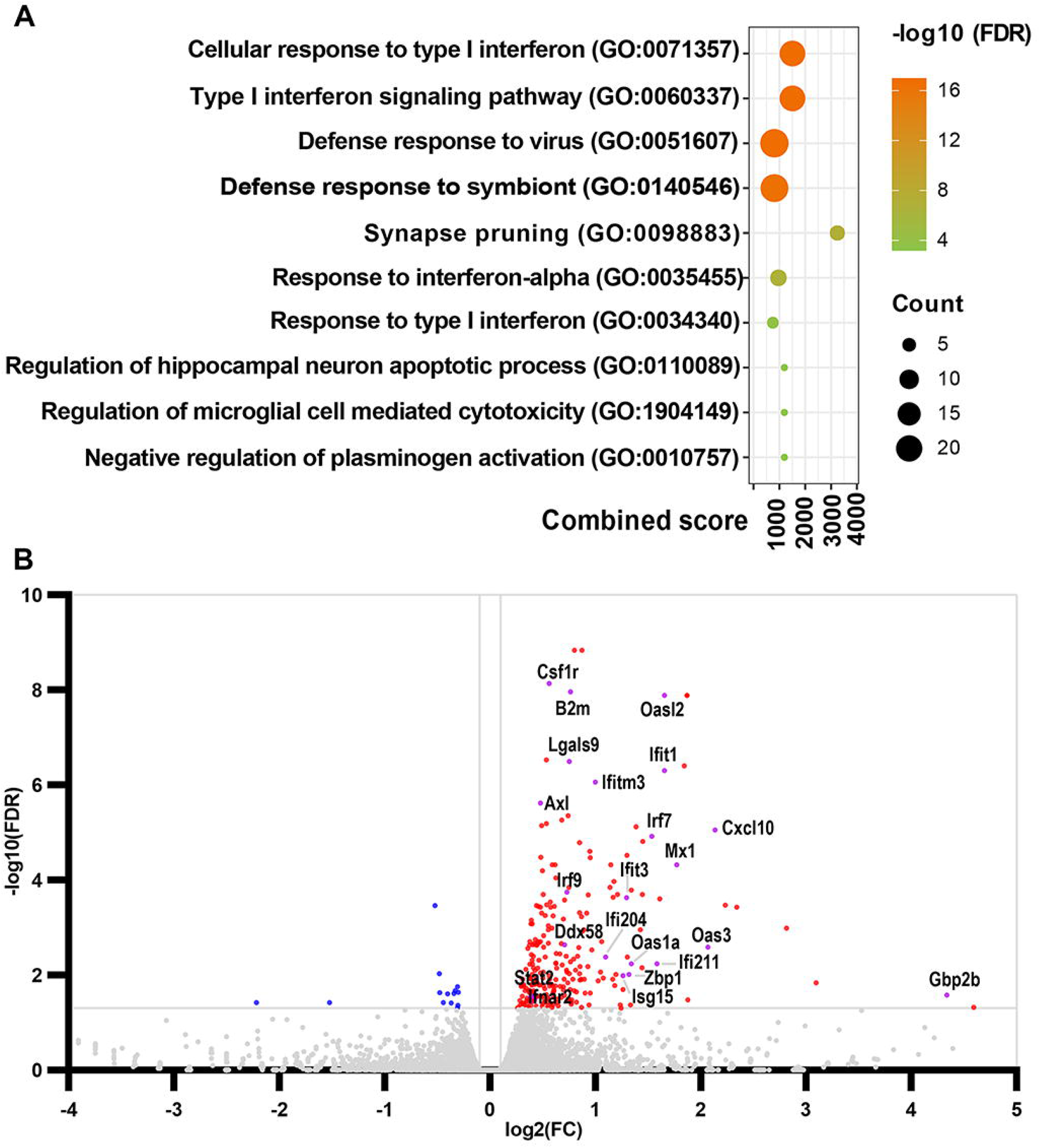
Effects of miR-34a KO on hippocampal transcriptome in Tg-SwDI mice. Bubble plot of GO enrichment analysis by Enrichr (A). The y-axis indicates the annotated terms of GO biological process in order of adjusted P value (-log10 (FDR)) and x-axis indicates combined score that represents a comprehensive metric combining the p-value and z-score. The bubble size represent the number of genes enriched in a specific GO term. The bubble color represents the adjusted P value. Volcano plot of DEGs between miR-34a^-/-^ and miR-34a^+/+^ Tg-SwDI mice (B). Red and blue dots represent upregulated and downregulated genes, respectively, and gray dots represent genes with no significant difference. The gray horizontal line represents 0.05 FDR and the gray vertical line indicate the log2FC threshold of 0.1.

### 3.5 MiR-34a knockdown enhances the type I interferon signaling pathway in BV2 cells stimulated with TLR7 agonist

To further investigate the role of miR-34a in type I interferon signaling, the mouse microglial cell line BV2 was used as an in vitro model. Gardiquimod, a well-characterized TLR7 agonist, was applied to assess TLR7-dependent activation of the type I interferon pathways in BV2 cells following miR-34a knockdown. Knockdown efficiency of miR-34a was confirmed by qPCR, showing a ∼160-fold reduction compared to the negative miR control (Figure 7A). Upon Gardiquimod stimulation, miR-34a-knockdown BV2 cells exhibited significantly increased expression levels of multiple type I interferon-stimulated genes (ISGs) and inflammatory markers, including Casp1, Ccl2, Ccl5, Cd70, Cd80, Cxcl10, Ifi204, Ifit1, Ifit2, Ifit3, Ifitm3, Il15, Isg15, Stat1, Oas1a, Oas1b, and Irf7 (Figure 7B, p < 0.05 or p < 0.01). These results suggest that miR-34a acts as a negative regulator of TLR7-induced type I interferon signaling.

**Figure 7.**
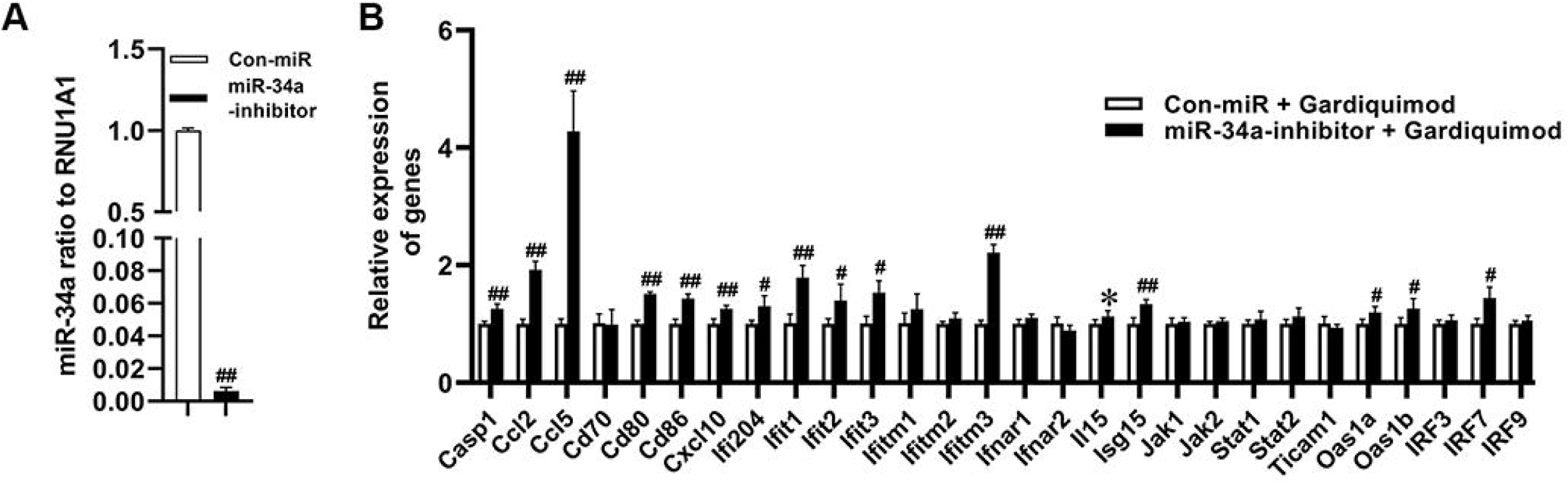
Effects of miR-34a knockdown on ISGs expression in gardiquimod-treated BV2 microglia. Depletion of miR-34a in BV2 cells by miR-34a inhibitor (A) and expression levels of the ISGs in miR-34a knockdown BV2 cells followed by gardiquimod treatments (B). (Data from three independent experiments, *P < 0.05, #P < 0.01 and ##P < 0.001).

## 4. Discussion

In this study, we demonstrate that miR-34a KO exhibits decreased anxiety-like behavior and improved memory in both non-transgenic and Tg-SwDI mice. MiR-34a KO further reduces anxiety levels specifically in Tg-SwDI mice. Moreover, miR-34a KO resulted in significantly elevated cerebral levels of soluble Aβ40, as well as insoluble Aβ40 and Aβ42 in Tg-SwDI mice, as measured by ELISA, compared to miR-34a WT Tg-SwDI mice. Based on these findings, we further analyzed activated microglia in the dentate gyrus and performed RNA-seq analysis on hippocampal tissues of miR-34a WT and KO mice, including both Tg-SwDI and non-transgenic groups. To our knowledge, these results are the first to show that miR-34a KO significantly increases activated microglia, along with upregulation of nucleic acid sensing and type I interferon signaling in Tg-SwDI mice.

Previous studies in the APP/PS1 AD mouse model have reported that miR-34a KO reduces amyloid deposition by downregulating β-secretase (not γ-secretase, as incorrectly stated in the abstract of the paper), and improves spatial learning and memory performance in the Morris water maze at 9 and 12 months of age (59). Cognitive improvement was associated with the upregulation of NMDA and AMPA receptors (60). In our study, we observed that miR-34a KO mice were more active and displayed reduced anxiety-like behaviors compared to miR-34a WT mice, regardless of transgene status. Additionally, miR-34a KO improved long-term spatial memory of mice, aligning in part with the prior findings. Consistently with previous studies (61, 62), the mild behavioral phenotype observed in Tg-SwDI mice, relative to non-transgenic controls, may be attributable to their very low expression levels of human APP and Aβ42 compared to other transgenic models that overexpress APP (63, 64). Despite this, Tg-SwDI mice recapitulate several key pathological features of AD and cerebral amyloid angiopathy (CAA), making them a valuable model for AD research (63). Because the non-transgenic mice also exhibited age-related cognitive decline in long-term spatial memory beginning at 8 months of age and beyond (64–66), our findings suggest that miR-34a KO mitigates both normal aging-associated memory impairments in non-transgenic mice and AD-like pathology-associated memory deficits in Tg-SwDI mice, potentially through distinct mechanisms.

Unlike the previous report (59), we found that miR-34a KO significantly increased the Aβ accumulation in the cerebral cortex of Tg-SwDI mice, compared to its controls. This increase included soluble Aβ40, as well as insoluble Aβ40 and Aβ42, while soluble Aβ42 remained undetectable in all groups. The increased Aβ levels were associated with increases in IFITM3 in the brains of miR-34a KO Tg-SwDI mice, compared to miR-34a WT Tg-SwDI mice. IFITM3 is a small transmembrane ISG and shown to enhance γ-secretase activity by binding to the γ-secretase complex (53). Therefore, the increased Aβ levels in miR-34a KO Tg-SwDI mice can be due to the increased expression levels of IFITM3. Interestingly, we found a decrease in the cortical Aβ42/Aβ40 ratio in miR-34a KO Tg-SwDI mice, compared to its controls, suggesting that Aβ40 increased more than Aβ42. It has been well established that Aβ42 exhibits more aggregation tendency and stronger neurotoxicity than Aβ40 (67). Additionally, it has been shown that 1) the increased ratio of Aβ42/Aβ40 promotes the formation of neurotoxic oligomeric assemblies in Aβ mixtures (51, 52), 2) small changes in this ratio markedly impact the characteristics of the Aβ mixtures, leading to synaptic damage and neurodegeneration (68), and 3) the relative ratio of Aβ peptides plays a more decisive role than the absolute amounts of Aβ peptides for the induction of neurotoxic conformations (68). These facts could be the reasons behind the improved memory in miR-34a KO Tg-SwDI mice despite the increases in Aβ load.

Several studies have demonstrated that miR-34a plays an anti-inflammatory role across various biological systems. In vitro, miR-34a mimics have been shown to reduce the expression of inflammatory molecules, including IL-1β, IL-6, CD11b, and iNOS, in LPS-stimulated BV2 microglial cells through suppression of the Notch1/Jagged1 signaling pathways (35). Additionally, miR-34a has been reported to protect the intestinal stem cell niche from inflammation-induced disruption in a model of Citrobacter-induced colon oncogenesis (69), further supporting its anti-inflammatory functions. Interestingly, previous studies using the miR-34a KO in TgAPP/PS1 mouse model did not report changes in microglial activation (59). This is possibly due to a ceiling effect, where microglia activation had already reached a plateau and could not be further elevated by miR-34a deficiency. As discussed earlier, Tg-SwDI mice express very low levels of human APP compared to other transgenic models, such as TgAPP/PS1 mice, which may make Tg-SwDI mice more sensitive to detecting modulatory effects on microglial activation (63, 64). The microglial cell body to cell size ratio, as measured by IBA1 immunostaining, has been positively correlated with microglial activation (54, 70). In the present study, we found that miR-34a KO significantly increased the microglial cell body-to-cell size ratio in Tg-SwDI mice based on morphometric analysis of IBA1-stained dentate gyrus sections, suggesting a suppressive role for miR-34a in microglial activation in this model.

While microglia were traditionally classified as pro-inflammatory (M1) or anti-inflammatory (M2), this dichotomy oversimplifies their diverse activation states. Single cell RNA-seq studies have identified distinct microglial phenotypes in neurodegeneration, including DAM, microglial neurodegenerative phenotype (MGnD) or activated response microglia (ARM) (71–74). Another microglial subpopulation that emerges not only in response to Aβ plaques but also during aging is defined as interferon-responsive microglia (IRM) that express several interferon-stimulated genes (ISGs) (73, 75). Importantly, dsRNA-induced IRM are essential for normal development of postnatal cortex and behavior, which is mediated by IRM that engulf excess neurons. Type I IFN deficiency during postnatal development in mice causes excitatory/inhibitory imbalance and tactile hypersensitivity (76). Homeostatic dysregulation of type I IFN signaling in microglia has been implicated in type I interferonopathies including Down syndrome (77). Triplication of Ifnar1, Ifnar2, and APP in patients with Down syndrome is thought to play a causal role in AD development. Accordingly, IRM appears to play a significant role in neuroplasticity under both homeostatic and diseased states. In this study, a comprehensive gene set enrichment analysis (Enrichr) on the data from hippocampal RNA-seq reveals that miR-34a KO activates IFN-I signaling, as shown by increases in nucleic acid sensors (Tlr3, Ifih1/Mda5, Ddx58/RIG-1, Ifi204, Ifi209, and Ifi211) and ISGs (Oas1a/b, Ifit3, Ifitm3, Ifnar2, Stat2, Irf7, and Irf9) in Tg-SwDI mice, but not in Non-Tg mice. Furthermore, miR-34a-inhibitor-treated BV2 microglial cells exhibited significantly increased expression of multiple ISGs found in IRM by gardiquimod, a TLR7 agonist, compared to the negative control, suggesting that loss of miR-34a sensitizes BV2 cells to TLR7 stimulation. Thus, our study using both in vivo (transgenic mice) and in vitro (cultured microglia) models suggests that miR-34a inhibit the transition of microglia to the IRM state. Interestingly, a recent study found that microglia increased expression of OAS1a together with other genes (Oas2/3, Mx1, Stat1/2, Ifit3, Ifitm3 and Usp18) in IFN signaling pathways and progressively gained an IRM state during aging in APPNL-G-F knock-in as well as wild-type C57BL/6J mice (78). APPNL-G-F knock-in mice express endogenous levels of APP with multiple mutations found in families with AD. Additionally, the same study found that the risk variants of OAS1 for AD and severe COVID-19 were associated with decreased OAS1 expression and exaggerated production of TNF-α with IFN-γ stimulation and that microglia with the IRM state in healthy humans increased during aging. The authors of the study suggested that the increased expression of ISGs with age might help reduce age-related damage by suppressing pro-inflammatory signaling (78). In line with this study, miR-34a KO significantly increased expression levels of Oas1a/2/3, Mx1, Stat2, Ifit3, Ifitm3 and Usp18 and improved spatial memory in Tg-SwDI mice.

In addition to upregulation of IRM signature genes, miR-34a KO induced significant increases in hippocampal mRNA expression levels of many DAM signature genes including Apoe, Tyrobp, B2m, Ctsb, Trem2, Cst7, Itgax, Clec7a, Axl, Csf1, and Gpnmb in Tg-SwDI mice (Figure 5A). DAMs are induced by and closely associated with neurodegenerative changes such as Aβ deposition (71, 72). Because miR-34a KO in Tg-SwDI mice increased levels of insoluble Aβ40 and Aβ42 in the hippocampus (Figure 3E) as well as the number of activated microglia in the dentate gyrus (Figure 4), increased expression of DAM signature genes in miR-34a KO Tg-SwDI mice can be due to increases in DAMs. Although DAM and IRM are distinct microglial activation states, the number of DAMs that express ISGs increases as β-amyloidosis in the brain progresses during aging of AD mouse models (5xFAD and APP-PS1 mice (79). Therefore, it is conceivable that miR-34a KO in Tg-SwDI mice may increase DAMs that express ISGs and/or may facilitate transition from DAM to IRM or vice versa. Further investigations by single-cell RNA-seq analysis are required to answer these questions.

In conclusion, we found that constitutive miR-34a KO altered microglial activation, decreased the ratio of cerebral cortex insoluble Aβ42 to Aβ40, enhanced memory, and increased expression levels of nucleic acid sensors and ISGs in Tg-SwDI mice. We showed that miR-34a knockdown strongly boosted microglial ISGs expression in response to a TLR7 ligand. Our study suggests that miR-34a inhibits the transition of microglia to the IRM state.

## Acknowledgements

We thank Austin Joseph Cyphersmith at Carl R. Woese Institute for Genomic Biology, University of Illinois Urbana-Champaign for his support with immunohistochemical imaging.

## Data availability statement

RNAseq files are available from the GEO database (accession number GSE235489).

