## Supplementary files for "MiR-34a deficiency enhances nucleic acid sensing and type I IFN signaling in a mouse model of Alzheimer’s disease"

**Supplementary Material**

**Supplementary Table 1**

DEG of experimental mouse groups

**Supplementary Table 2.**

GO_BP from Enrichr analysis

**Supplementary Table 3.**

IRM signature genes significantly upregulated in miR-34a-deficient Tg-SwDI mice and potential miR-34a target genes.
